# Does internal metabolic state determine our motor coordination strategy?

**DOI:** 10.1101/312454

**Authors:** Scott V. Taylor, A. Aldo Faisal

## Abstract

Motor coordination requires the orchestration of multiple degrees of freedom in order to perform actions. Humans display characteristic and predictable reaching trajectories even though multiple trajectories are possible. Computational theories of motor control can explain these reaching trajectories by assuming that subjects orchestrate their movements to minimise a cost function, such as end-point variability or movement effort. However, how internal metabolic states influence decision making and sensorimotor control is not well understood. Here we measure human behaviour during a centre out reaching task in two distinct metabolic conditions in the morning – after having had breakfast and not. We find that humans alter their patterns of motor coordination according to their internal metabolic state and that this change in behaviour results in a 20% lower task-related energy expenditure when fasted. We suggest that movements are orchestrated according to different criteria in different metabolic states so that metabolic costs are reduced in low metabolic states. We also predict that motor coordination strategies take the metabolic costs of specific muscle groups into account when planning and executing movements. Thus, metabolic state may alter the computational strategies of decision making between animal-based experiments (in typically low metabolic conditions) and human psychophysics experiments (in typically high metabolic conditions).

## Introduction

The brain is thought to behave in a Bayes-optimal manner ^1–3^, including the way it controls the motor system ^4^. Bayes-optimal behaviour defines the optimal behaviour in the presence of uncertainty and maximises the probability of a successful outcome given knowledge of a task and one’s prior beliefs about the world. For example, human behaviour in a ball-catching task can be explained as a near optimal combination of sensory and motor noise ^5^. It has been suggested that evolution gives rise to optimal behaviours by selecting for behaviours that maximise survival ^6^. One key feature of motor behaviour is coordination, where multiple degrees of freedom, such as muscles, joints and limbs ^7^, are involved in a task.

Optimal feedback control theory (OFCT) has been very successful in explaining human motor coordination in a principled manner ^8,9^ and has been shown to predict a large variety of movement data ^8–11^. For example, OFCT predicts that we reduce variability in task-relevant dimensions and allow it to increase in task-irrelevant ones, thus improving task performance^11^.

OFCT derives motor control strategies from the minimization of a cost function that is dependent on motor commands, state variables and task variables ^12^. State variables describe the current state of the dynamical system and commonly include the current position and velocity of the system ^9,13–15^. Task variables describe the objective of the task and commonly include a target position that should be reached by the system ^9,13^.

There are a number of cost functions that have been proposed in literature, which can be divided into intrinsic costs and task-based costs. Intrinsic costs are costly in and of themselves, rather than due to an external factor. Often they describe the usage of a limited resource that may be difficult to recover. The motor system has many examples of intrinsic costs within its function, for example, energy usage, motor variability and fatigue. Certain solutions to a task may be considered more desirable if they are associated with lower intrinsic costs yet reducing these costs may not improve task performance and may even decrease task performance. These costs do not directly arise from the property of a task hence they should apply widely and be conserved over many diverse motor tasks.

Task-based costs arise from the properties of a motor task. For example, when defining costs for a targeted reaching task, there would be a certain cost on missing the target, a cost for reaching the target outside of a strict time window and also a cost on final velocity such that you would need to come to a complete stop at the target. These costs define the task, and similar costs can be applied to many tasks, although they do not generalise well between tasks. Task-based costs are necessary in the field of motor control for formalising the underlying problem such that control algorithms (such as optimal control) can be applied to them.

When making decisions, the internal state of animals, including humans, changes the weighting between their different options. Metabolic state, which encompasses the available energy levels within the body including energy stores such as fat cells and blood glucose, is one of the internal states that may be important to the human motor system. Efficient and sustainable use of energy is of vital importance to organisms ^16^ and their behaviour. Movement accounts for up to 70% of daily energy usage ^17^ and therefore energy sets a fundamental constraint for motor behaviour. The brain may thus have evolved to plan and control movements in a sustainable manner. Therefore, the amount of energy available to act – our internal metabolic state – may be an important factor used by the brain for motor planning and coordination. While it seems intuitively obvious that we would choose to move more slowly, or not at all, when we have little energy, we may not have the option for slower movements or inaction in many real-world situations. This is the case for motor coordination tasks such as reaching, where the motor system has to choose a suitable movement trajectory from the theoretically infinite ways to reach an end-point, within a given time frame. It has been suggested that the brain may choose these movement strategies so as to minimise the impact of muscle force variability ^18^, effort ^19^ or jerkiness of movements^20^.

It has previously been shown that subjects’ movement trajectories in reaching tasks can be explained by a choice of joint movement that minimizes the influence of signal-dependent motor noise in muscles, thus adapting a reaching strategy that minimizes end-point variability ^18,21^. Minimisation of motor noise can also explain more complicated motor tasks, such as manipulating a non-rigid object with an internal degree of freedom ^22^. However, O’Sullivan ^19^ showed that a finger force production task behaviour is better explained by minimum effort rather than minimum variability strategy, with a more even force distribution across fingers than would be predicted by the measured variability. This suggests that the motor system may trade off different motor costs against each other. More recently, Huang ^23^ showed that practice, or motor learning, reduces the metabolic cost of reaching movements in force fields, revealing that metabolic efficiency can be improved by the brain.

In this study, we ask whether the internal state of the human body influences decision making by the brain, that is, whether the internal metabolic state can affect decisions about motor coordination if cost functions remain the same. We hypothesize that, when confronted with motor coordination problems with fixed time constraints, our brain may exploit knowledge about its internal metabolic state to inform its motor system. For example, our brain may prioritise more energy-efficient motor coordination strategies when low on energy over other strategies that promote desirable features such as movement accuracy, which may be chosen when high on energy. We directly test this using a centre-out reaching task for subjects in two metabolic states, high (fed) and low (fasted), while keeping movement time and thus movement velocity constant. By analysing the reaching trajectories, we can observe task-specific motor coordination strategies and performance, which allows us to analyse the decision-making processes in the brain. Observing changes in these measurements between metabolic states allows us to determine whether or not these decision-making processes depend on internal metabolic state.

## Results

In order to test if a subject’s internal metabolic state has an impact on the strategy for motor coordination, we asked eight subjects (aged 23-29) to perform a centre-out reaching task in two distinct metabolic states and measured the position of their hand throughout the movements. We combined this with indirect calorimetry to estimate energy expenditure during the task using a Quark RMR metabolic cart.

During reaching movements, the metabolic cost was significantly higher (Student’s t-test, p = 0.000067) than during the resting phases in both high and low metabolic states (Fig. 1), as expected. Interestingly, metabolic energy expenditure was significantly elevated (Student’s t-test, p = 0.000036) in both experimental phases when fed compared to fasted (Fig. 1). In order to take the different resting energy expenditure for the two metabolic states into account, we measured task-related energy expenditure, and found the difference in metabolic expenditure for reaching minus rest to be significantly larger when fed versus fasted (Student’s t-test; p=0.012, 2.4). Indeed, task-related energy expenditure increased significantly in 7 out of the 8 subjects (Student’s t-test, p = 0.0058, 0.000020, 0.00075, 4.0e-14, 0.0026, 3.6e-7, 4.6e-10, 1.1e-10). When comparing task-related energy expenditure for the first versus the second experimental session, there was no significant difference within the group (Student’s t-test; p = 0.15).

**Figure 1:**
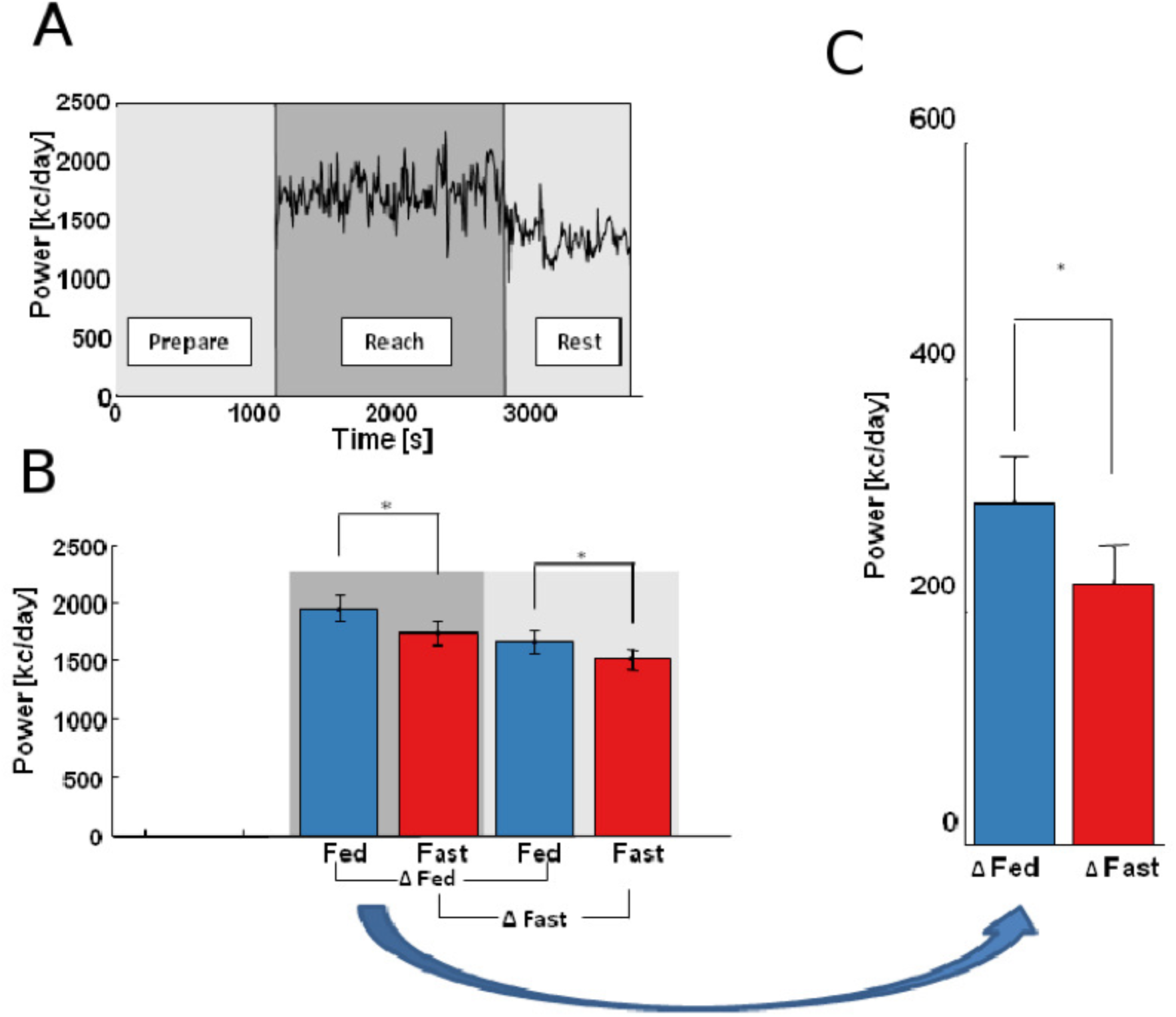
Task-related energy expenditure depends on internal metabolic state. **(a)** An exemplar raw metabolic expenditure trace. **(b)** Metabolic energy expenditure was lower during the rest phase (light shading) compared to the reach phase (dark shading) and when subjects were fasted (red) compared to fed (blue). **(c)** Task-related metabolic energy expenditure was reduced when fasted.

To confirm whether our metabolic regime resulted in a measurable difference in internal metabolic state, we measured the respiratory exchange ratio (RER) of subjects, and found it was significantly lower when fasted for 7 out of 8 subjects (Student’s t-test for each subject: p = 3.4e-61, 2.2e-121, 1.4e-176, 7.2e-149, 7.0e-183, 5.5e-124, 1.5e-10) and for the group (Student’s t-test; p = 0.0075). The significant change in RER confirmed that the majority of subjects fasted as instructed (see Methods) and that the fasting regime produced a measurable change in metabolic state. We can thus conclude that internal metabolic state affects task-related energy expenditure during this motor task.

We hypothesised that reduced task-related energy expenditure in the fasted state might be due to trajectory changes during the reaching task and therefore analysed the subjects’ hand reaching trajectories. Reaching trajectories were characteristically curved in each direction (Fig. 2a-d), although there were clear differences between subjects. Importantly, reaching trajectories were significantly different between metabolic conditions for 7 of the 8 subjects (non-parametric bootstrap with Bonferroni correction over directions; p = 7.7e-8, 0.00059, 0.00049, 0.078, 0.045, 0.000014, 0.025, 0.0076). For the group of subjects, task-constrained positions (start and end points) did not change significantly (Student’s t-test, N = 8, p = 0.13). However, reaching trajectory mid-points showed significant differences (Student’s t-test, N = 8, p = 0.020, Fig. 2b), shifting to the right when subjects were fasted (Fig. 2e, f). This change was also evident even if we factored in on a trial-by-trial bases the negligible differences in start and end positions (Student’s t-test, N=8, p = 0.027). Reaching trajectory mid-points were not correlated with start (Spearman’s correlation test, → = 0.087, p = 0.84) or end positions (Spearman’s correlation test, → = 0.26, p = 0.52). We thus conclude that despite the large difference between subjects’ reaching trajectories, all subjects change their trajectories in a similar way between metabolic states.

**Figure 2:**
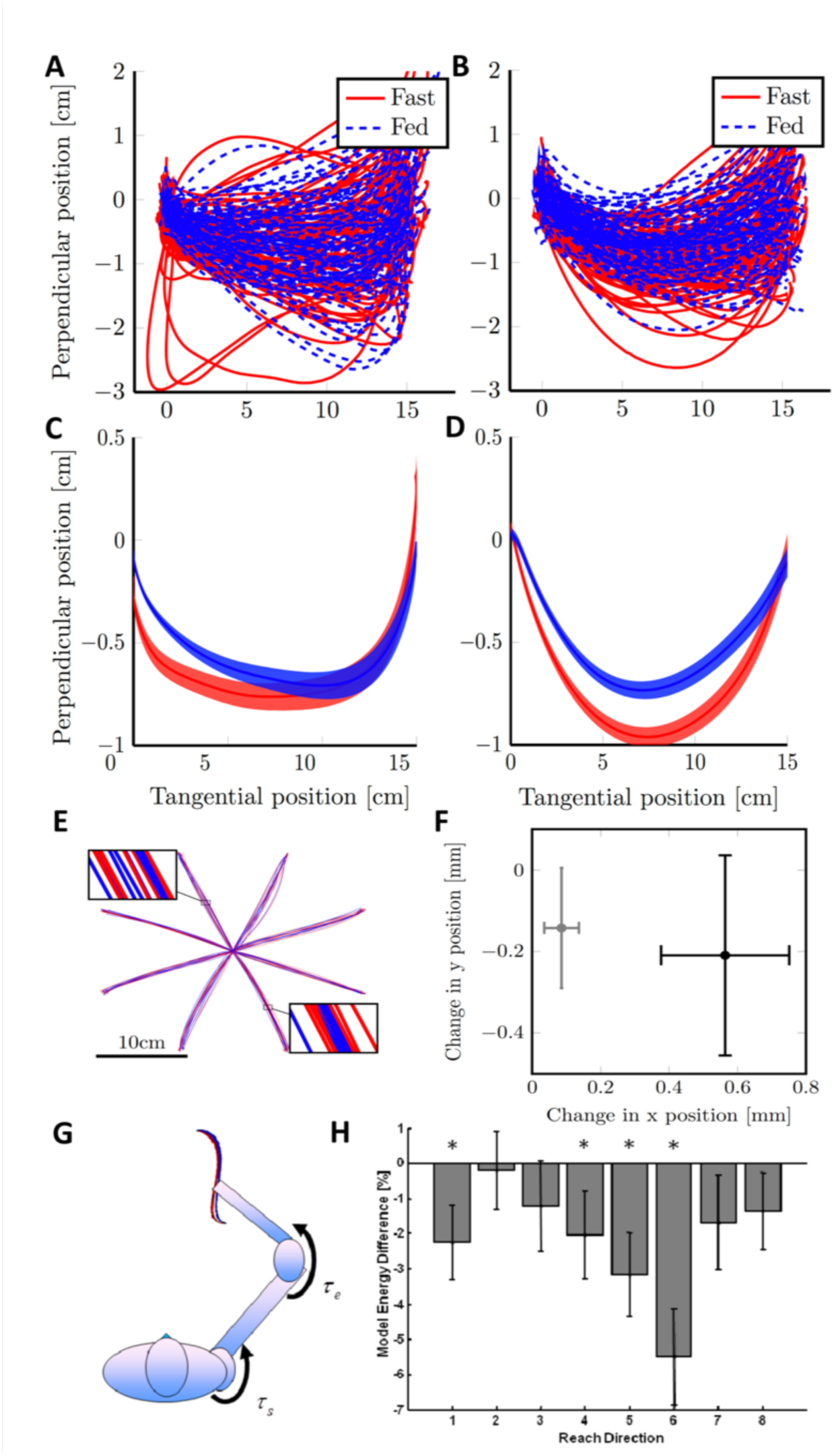
Reduced energy expenditure during fasting associated with change in reaching trajectory. (**a**) and (**b**) Raw reaching trajectories in one direction for two exemplar subjects in fasted (red) and fed (blue) metabolic states. (**c**) and (**d**) Mean reaching trajectories for the same two subjects in the same direction with shaded areas denoting standard error of the mean. Note the curved shape of mean reaching trajectories in both fed and fasted metabolic states. The coordinate frame was defined by a line from the central point to the target, with one axis tangential to this line and the other perpendicular to it. Due to the inherent geometry of the task, the perpendicular dimension is displayed as a different scale to the tangential dimension. (**e**) Mean Reaching Trajectories for all subjects in all reaching directions for both fed and fasted metabolic states. Mean reaching trajectories displayed subtle curvature in several directions. Insets show two reaching midpoints on a larger scale to highlight the differences in reaching trajectories between metabolic states. (**f**) Mean ± standard error change in reaching midpoint (black) and task-constrained positions (grey) across subjects. Note, the significant shift of reaching trajectory midpoint to the right (x-direction) when fasted (*p* = 0.020). Task-constrained positions moved slightly to the right, but this shift was not significant (*p* = 0.13). (**g**) Diagram of the 2-Joint arm model denoting torques on each joint and dynamic equation. (**h**) Energy difference between performing reaching trajectories in each experimental metabolic condition. The error bars include +/− standard error of the mean due to experimental variability and ±1% model parameter variation. Note, that all directions decreased energy expenditure as a group and individually also directions 1, 4, 5 and 6 were significantly less costly (non-parametric bootstrap, *p* = 0.015, 0.045, 0.003, 0.001) denoted by *.

An important premise of our task was that subjects would move at the same speed in both metabolic conditions. We therefore gave strict negative feedback to any reaching movements that were completed in a time that fell outside of a tight time window during the experiment. To verify if this was sufficient to maintain movement speed, we checked if the rate at which subjects performed reaching movements was successfully controlled between metabolic conditions. Inter-trial times did not significantly change (Student’s t-test, p=0.64) and group mean peak velocities were conserved (Student’s t-test, p = 0.276). The rate at which subjects performed reaching movements was thus successfully controlled between metabolic conditions and any changes in energy expenditure cannot be due to subjects moving more slowly.

In order to determine if the change in reaching trajectories might have caused the observed decrease in energy expenditure, we used a simple arm model (Fig. 2g) to measure mechanical work. We tested each subject individually and found the majority (6 of 8) of subjects’ mean reaching trajectories were associated with lower levels of mechanical work when fasted (Student’s t-test, p = 1.3e-35, 4.7e-27, 0.0048, 6.8e-32, 3.0e-20, 5.7e-5), as were the mean trajectories of the subjects in the fasted state (Student’s t-test, p = 0.0061) as well as for individual directions of reaching (see Fig. 2h). The mean reduction in mechanical energy expended by the model in the fasted state was 3% lower that the expenditure when fed. This indicates that the change in reaching trajectories was responsible for at least some of the decrease in energy expenditure

To test if there was a trade-off between energy expenditure and variability in the task, we looked for a negative correlation between the change in metabolic energy expenditure with metabolic state and the change in reaching error with metabolic state for each subject. Subjects that tended to reach closer to the target when well fed compared to when fasted reached it in a less metabolically efficient manner when fed. However, while the change in energy expenditure was significant between metabolic states, the change in end point error was not (Student’s t-test, p = 0.27). Indeed, three subjects using slightly more energy when fed also increased their end point error. This may have been due, in part, to subjects being rewarded or penalised for stopping within or outside the target rather than penalised based on RMS error to the centre of the target. Despite this, we found a significant correlation (Spearman’s correlation test; p = 0.048) between the difference in end point error and the difference in task related energy expenditure (Fig. 3). Energy and variability were thus negatively correlated, suggesting that as subjects chose to alter their motor coordination when fasted, they did so at the expense of greater task accuracy.

**Figure 3:**
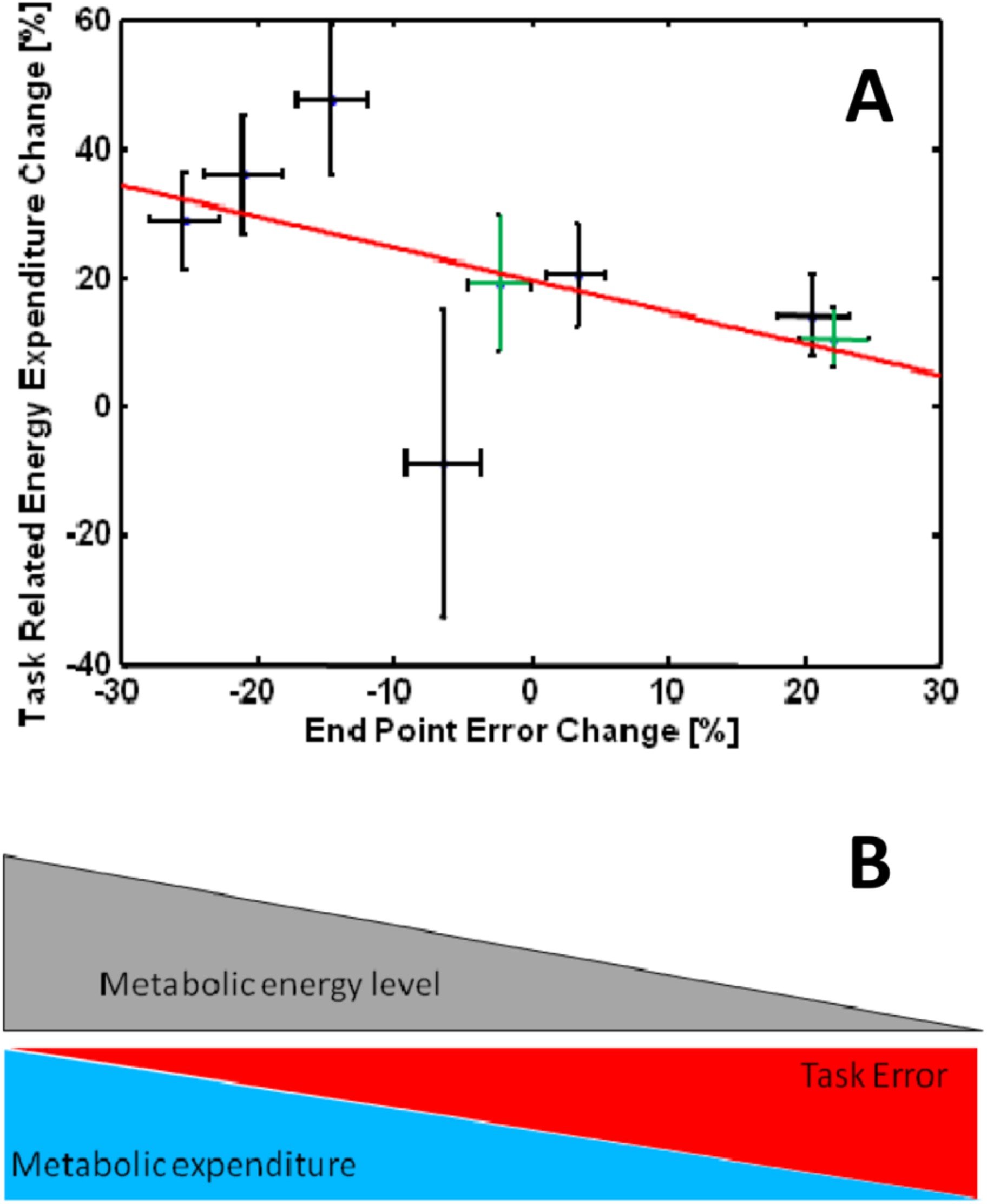
Task accuracy is impaired during energy conservation when fasted (**a**) Change in task-related energy expenditure plotted against the change in end-point error between experimental metabolic conditions, for each subject. Subjects with a positive task-related energy expenditure change used more energy when fed. The three subjects with the largest percentage change in energy expenditure used more energy in the second session and had the largest percentage improvement (decrease) in end point error. The two subjects with the smallest percentage positive change in energy expenditure had the largest percentage degradation (increase) in end point error. This trend lead to a significant negative correlation between percentage change in energy expenditure and percentage change in end point error (Spearman’s correlation test; *p* = 0.048). One subject did not show a significant change in energy expenditure, leading to a much larger uncertainty in percentage energy expenditure change compared to the other subjects. Error bars denote standard error of the mean. Subjects that fasted for the second session (green) and subjects that fasted in the first session (black) were not significantly different (Student’s t-test, p=0.28, p=0.59) in either dimension. (**b**) A diagrammatic representation of a suggested trade-off that could cause the effect seen in (**a**). As metabolic state decreases, energy efficiency is considered more important, and task error is considered less so. This is predicted to lead to a correlation between increased energy efficiency and increased task error, as observed in this study.

In conclusion, we have shown that motor coordination changes with metabolic state such that there is an increased mechanical energy efficiency of reaching movements when fasted and an associated decrease in task-related energy expenditure. We also observed a correlation between increases in energy efficiency and increases in task error suggesting a trade-off between two undesirable aspects of motor coordination tasks: energy and variability.

## Discussion

We have demonstrated that internal metabolic state plays a determining role in the motor co-ordination of arm reaching movements. Specifically, arm reaching movements are less metabolically costly when fasted.

Task related energy expenditure was significantly reduced when fasted. By controlling for resting energy expenditure, we rule out that this difference is due to digestive processes, and postural muscles which are equally active during the task and the resting phase. Central Nervous System and cardiovascular expenditure may be raised due to the activity required to perform the task yet there appears to be no clear reason why energy expenditure from these processes alone would be significantly higher when in a high internal metabolic state. The remaining component of energy expenditure, mechanical work, is the only component that was not present at all during the resting phase and was expected to account for the majority of the task-related energy expenditure.

Depending on metabolic state, movement strategies differ. Given the redundant nature of our motor system, when confronted with the same movement task we change our motor coordination based on our metabolic state. This is possible due to the highly redundant nature of the human motor system, i.e. we can reach to the same point via many different paths. While there were many differences in reaching trajectories between subjects, changes between metabolic states were conserved between subjects. This suggests that once controlling for differences between subjects, be they habitual or due to differences in mechanical properties such as arm segment lengths or muscle sizes, changes in metabolic state produce similar changes in reaching trajectories.

We found that the mechanical energy expended during reaching was lower when fasted in a simple arm model. Simple two-joint arm models are regularly used in motor control studies ^13,20,24^ as they are thought to capture many important aspects of the mechanics of the human arm. This suggests that some of the changes in task related energy expenditure are caused by the changes in motor coordination and that subjects moved towards a more metabolically efficient motor coordination strategy when fasted. Subjects respiratory quotient was measured as an indication of internal metabolic state. A higher respiratory quotient is associated with the burning of carbohydrates versus lipids ^25^. As humans tend to preferentially consume carbohydrates over lipids ^26^, respiratory quotient tends to decrease during fasting ^27,28^. A single subject’s respiratory quotient was increased during the fasted session, which could be explained by either the subject fasting before both sessions or the subject consuming an overwhelmingly fatty meal before the fed session. The lower respiratory quotient observed during the fasting state shows that the majority of our subjects did fast and that our fasting regime produces distinct internal metabolic states.

Respiratory quotient is a better measure of internal metabolic state than for example blood glucose levels. The circadian rhythm and insulin can combine to produce wide variation in measurements of blood glucose after a morning meal, potentially as low as 65 mg/dl ^29^, depending on meal composition and time of eating ^29,30^. A long fast of 52 hours can be expected to reduce blood glucose levels from 88 mg/dl to 66 mg/dl ^31^, so our much shorter period of fasting is expected to have a smaller effect on blood glucose levels. This is important in and of itself, as a significant reduction in blood glucose may impact on muscle function and we wish to avoid this. Hence, for subjects following our metabolic regimen we may not have been able to discriminate between fed and fast metabolic states using measurements of blood glucose.

We could ask why humans do not always use the most metabolically efficient solution to a motor task. We have found that subjects’ increase in error in the task correlates with their increase in metabolic efficiency, suggesting that there is a fundamental trade-off occurring, with the balance between task accuracy and metabolic cost determined by internal metabolic state. When calculating correlation between the change in task related energy expenditure and the change in end point error, we assumed that there was no difference between subjects that fasted first and subjects that fed first. While we could not reject this hypothesis, the low sample size (N=8) does not provide strong evidence.

The change in task related energy expenditure (approximately a 20% reduction) did not match well the change in mechanical energy expenditure in our model (approximately a 3% reduction). This may be due to problems with the assumptions underlying the model such as assuming that torque is always produced with 100% efficiency. Because our muscles are comprised of muscle fibres with differing metabolic properties ^32^, the results obtained in this study could be explained by a change in emphasis from activating metabolically more costly muscles (or muscle fibres) to more efficient muscles (or muscle fibres) as internal metabolic state decreases.

An alternative explanation for the results of this study are that subjects change their behaviour due to slow learning processes. While we have not fully controlled for the effects of motor learning in this study, alone it cannot explain the results presented here. More of our subjects fasted in the first session which produced conflicting learning and metabolic state hypotheses. The learning hypothesis was that metabolic expenditure would be lower during the second session as it has previously been found that metabolic expenditure decreases with motor learning ^23^. The metabolic hypothesis was that metabolic expenditure would be lower when fasted as the motor system was predicted to utilise motor redundancy to perform the task in a more energy efficient manner. We observed an increase in metabolic expenditure during the second session and a decrease in metabolic expenditure when fasted. This indicates that the effects of internal metabolic state were dominant within our experimental paradigm. The increased metabolic efficiency observed in this study can also be linked to studies of animal metabolic state and activity levels. In this study, our fasting regime should have reduced the level of leptin in the blood ^33^ and we observed an increase in metabolic efficiency of movement. In animal studies, the leptin pathways can be interfered with using viral vectors or injecting chemical antagonists which results in reduced activity ^34,35^. The same mechanisms of the brain that results in a higher probability of inactivity in animals may also seek to reduce the amount of energy expended during an activity in humans.

Our study shows that the human motor system produces more energy efficient movements when in a low metabolic state. This could be a big problem when attempting to lose weight solely through calorie restriction. If reducing calorie intake produces a low internal metabolic state, routine motor tasks may be executed in a more efficient manner. This may lead to a rebalancing of energy intake and expenditure and could prevent weight loss.

This study points the way to novel avenues for tackling obesity, a key problem in the Western world ^26^. If the mechanisms that inform the motor system of internal metabolic state can be identified and altered, the motor system could be encouraged to produce less metabolically efficient coordination strategies. This would in turn lead to an increase in energy expenditure with no need for a change in routine.

We have shown that human motor coordination can be determined by non-neuronal physiological states. In order to make accurate predictions of human performance in a task without fitting a model with large numbers of free parameters, it is necessary to understand what quantities the brain cares about and how these affect the resultant behaviour. There may also be other internal states that modulate human motor coordination. By investigating the effects of these on motor coordination we will develop a greater understanding of the decision-making processes in the brain.

Traditionally, decision-making is often thought of as a conscious selection of a choice from a range of options. Here, we talk about decision making in the motor system, which is not to suggest that the motor system is making conscious choices, but rather to take the view that decisions are a more general process where one of several actions is taken in the presence of uncertainty of outcome based on the potential value of each action. In many simple motor tasks, the human motor system can perform a number of different actions, yet often actions are chosen in a characteristic manner. Given the existence of fundamental motor costs such as energy usage and motor variability, the probable outcome of each action can be valued relative to the probable outcomes of other actions. Thus, motor coordination tasks can be thought of as continuous decision-making processes^36^. Even if the outcome is determined by hardwired responses, a decision has still been made.

A promising direction of future study will be to investigate other determinants of motor coordination strategy and produce a more detailed model of how the brain evaluates non-neuronal physiological states and uses them to inform decisions about motor control. This will lead to improved predictions and understanding of characteristic human motor behaviours, aiding in the diagnosis and mitigation of motor disorders and improving the quality of life of those that suffer from them.

Our study shows that it is important to consider metabolic state when performing motor control experiments. If metabolic state is not controlled for, this significant source of variability in behaviour may bias the results of a study. Many animal studies use food or liquid (juice) rewards with calorie content as a source of motivation to perform a task well. The need to supply a clear reward signal and thus potentially affect metabolic states in the many tasks using animal behaviour reporting may thus engage and or shift neural computations and decisions into a narrow response regime. It may be advisable therefore to consider metabolic state as a potential complicating factor in these studies.

Each of our subjects followed an unknown diet before each experimental session. We were able to verify that the respiratory quotient changed due to the breakfast/no breakfast protocol. The change in respiratory quotient was consistent with a reduction in available energy for all but one subject but it is unclear if the variability in this effect was due to differing responses from the subjects’ metabolism or due to differing levels of calorie restriction. The level of calorie restriction that had been undergone is unlikely to have been the same for each subject and the logistics required to quantify it were prohibitive. Thus, there could well have been a great variability in the number of calories consumed by the experimental subjects leading up to each experimental session.

Much more can be done with regards to controlling calorie intake and restrictions in animal studies. For example, in one recent animal study, blowflies were deprived of food for 3 days^37^ which allowed successive decreases in neural activity related to the visual system to be plotted over time. This level of food deprivation is clearly not ethical with human subjects as the risk to health, the practicalities of monitoring and enforcing such a regime. On the other hand, it is often impractical to perform precision motor coordination experiments with animal subjects due to difficulties in motivating and teaching animals to perform the tasks.

## Methods

### Subjects

Healthy right-handed subjects (N=8; aged 23-29) participated in the study. The experiments were carried out in accordance with institutional guidelines, and a local ethics committee approved the experimental protocols.

### Protocol

Subjects were sat in a Virtual Reality Rig (Fig 4.A-B), where visual feedback was projected into the plane of movement via a mirror system. The position of each subject’s right hand was recorded using a Liberty magnetic tracking system (Polhemus; Colchester, VT, USA) and the weight of their arm was supported by an air puck which allowed frictionless movement within a horizontal plane. A sports-car driving seat constrained movement of the torso so that movements of the arm were not confounded with upper body movements. The positioning of the seat was constrained by a tight fitting semi-circular cutout in the surrounding table and subject’s feet were supported by an adjustable footrest. Lighting conditions were carefully controlled using black-out curtains to remove natural light. Subjects’ wrists were constrained with a sports support to discourage large movements of that joint.

**Figure 4.**
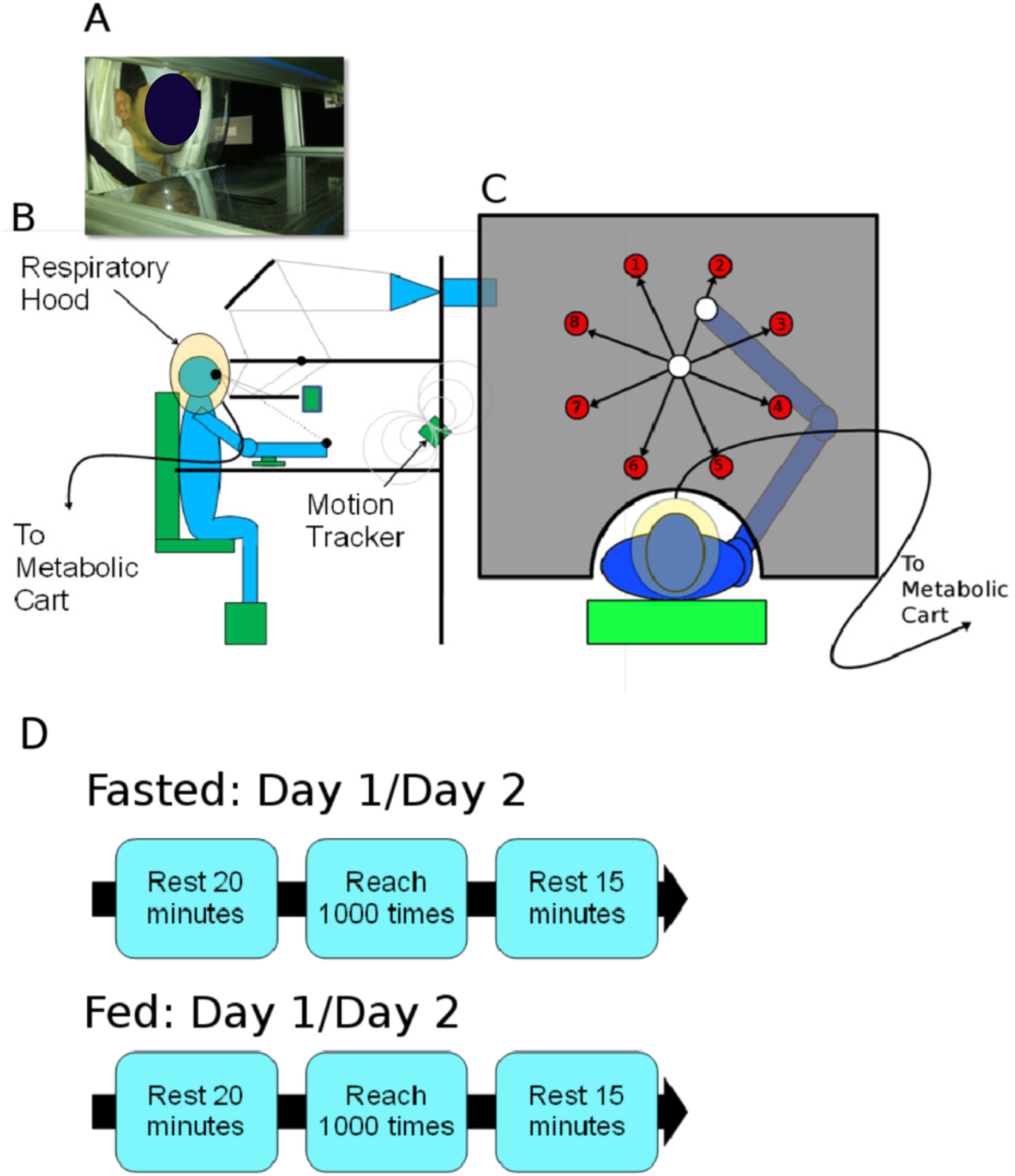
**A**. A subject sits in the setup ready for the experiment. A metabolic cart measured expired respiratory gases that were captured by a ventilated hood. **B**. Subjects sat in a sports-car driving seat with their arm supported on a frictionless surface. A motion tracker recorded the position of their right hand. Visual feedback was projected via a mirror system. **C**. Subjects performed centre-out reaches from the white sphere to one of eight targets positioned as shown. **D** Subjects followed a strict metabolic protocol during the experiment. A pre-task resting period was used before each session to reduce any metabolic effects of activity prior to the experimental sessions. A post-task resting period was used to measure task-irrelevant energy expenditure.

Subjects were asked to perform centre-out reaches with their right hand to one of 8 targets within a horizontal plane (Fig. 4.C) in random order with a 1.12kg weight strapped to the wrist. There were 1000 randomly ordered reaches per experimental session. Subjects were allowed to familiarise themselves with the setup prior to each experimental session. The centre out reaching task began when participants moved their hand (position was indicated with a sphere of radius 1cm) to the visual workspace centre (approximately 30cm in front of the torso and central with respect to the shoulders); then a 1.5cm radius target sphere appeared 15cm away in one of 8 equally spaced directions.

Subjects chose when to start reaching towards the target, having to enter a region of radius 2cm centred on the target sphere within 0.33s. After a further 0.25s, if the subject’s position was within the target sphere, the subject would be informed of a successful trial; otherwise the subject would be informed of a mistrial due to missing the target. If the subject reached the region around the target sphere before 0.24s, the subject would be informed of a mistrial due to reaching too quickly and if they had not reached the region within 0.33s the subject would be informed of a mistrial due to reaching too slowly. At all times subjects could see the position of their hand, upon reaching the target it changed colour from red to yellow. Feedback was given in the form of a score that increased by 1 for successful trials and decreased by 1 for unsuccessful trials. Participants also received visual feedback if they were too fast, too slow or if they missed the target. If the movement did not enter an area larger than the displayed target (an area of radius 2cm centred on the target), or if the movement was deemed too fast or too slow the trial was excluded from the analysis of reaching trajectories. The first 10 minutes of data from each day’s reaching phase were removed so as to better measure the steady state values as opposed to transient responses (e.g. potential effects of learning/warming-up).

Written instructions were displayed at the top of the workspace and were verbally reiterated by the experimenter. A score was displayed in the top-right corner of the screen as an integer value.

### Metabolic Conditions

Each subject performed the experiment on two mornings starting between 9am and 10.30am, on the first morning they were either in a fed or a fasting state and on the second fasted or fed. The fasting state was defined as not eating after 8pm the preceding evening and skipping breakfast on the day of the experiment. N=6 subjects fasted for the first session and N=2 subjects fasted for the second session. Each experimental session (Fig. 4.D) began with 20 minutes of preparatory rest, during which the subject was sat in the experimental rig but was instructed not to move. This phase was designed to wash out the metabolic effects of any prior activity. The subject was then instructed to begin the centre-out reaching task which continued for 1000 trials or approximately 30-40 minutes.

Finally, the subject was again instructed to rest for 15 minutes in order to measure task-irrelevant metabolic energy expenditure including basal, digestive and postural expenditure. The first 5 minutes of data from the final rest phase were removed so as to allow the metabolic data to settle down to steady state resting levels.

Each subject wore a Ventilated Hood which collected expired gases. Ventilated Hoods have been shown to be more sensitive than conventional Mouthpiece and Nose-Clip systems ^38^. We used a Quark RMR metabolic cart (Cosmed; Rome, Italy) to measure Oxygen consumed and Carbon Dioxide produced. From these, we calculated energy expenditure and RER. Energy expenditure (including expenditure related to digestive processes, postural muscles, cardiovascular, CNS and mechanical work) was calculated using the modified Weir equation ^39,40^. The device was calibrated before every use with a certified gas mixture.

Task relevant energy expenditure was calculated as the difference between metabolic expenditure during the task phase and the final resting phase. A non-parametric bootstrapping test was used to test for a difference in task-relevant energy expenditure between fasting and fed experimental sessions. RER was calculated as the ratio of Oxygen consumed versus Carbon Dioxide produced and we similarly tested for a difference in RER between fasting and fed experimental sessions.

### Statistical Analysis

A distribution of subjects’ mean reaching trajectories were calculated with non-parametric bootstrapping in order to compare fasting and fed motor coordination strategies. By integrating the area between two randomly selected traces, a test statistic could be calculated for the chance that there was a difference between the fed and fasted reaching trajectories. The expected mean area is zero as there is equal change of a negative area as a positive area. A mean area that is significantly different from zero demonstrates significant difference between two populations of reaching traces (fig. 2.3).

To test for changes in the group mean trajectories we examined the mean difference in reaching midpoint position as this was the least constrained by the task. We also examined this difference after controlling for changes in task-constrained positions (positions at the start and end of each reaching movement).

A non-parametric Spearman’s rho was calculated to test for correlation between the difference in end point error (shortest distance to the target during a reaching movement) and the difference in metabolic energy expenditure, both differences calculated between fasted and fed cases.

### Mechanical Modelling

We used a 2D two joint arm model to calculate the mechanical work needed to produce the group mean movement trajectories in each of the 8 movement directions observed during the fed and fasted regimes. We did this by calculating the inverse dynamics of the arm, allowing us to calculate mechanical work as the product of torque and angular velocity.

Our arm model has arm segment lengths of 30cm and 34cm, segment masses of 1.4kg and 1.1kg and a 1.1kg weight located at the wrist. The joints are subjected to 0.5kgm2s-1 of friction. The shoulder of the model is located in a suitable position relative to our experimental setup. The model follows the following dynamics:

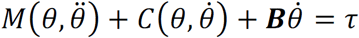

Where C is Coriolis and Centripetal forces, B is the joint friction matrix and M is the inertia matrix. The torque τ can be calculated by supplying an experimental movement trace. Joint angles are calculated from the Cartesian coordinates of each sensor reading then differentiated twice to calculate the two time derivatives. The above equation can then be solved for torque, τ. Mechanical energy, E, is then defined as:

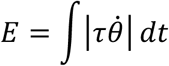

Changes in mechanical energy expenditure derived from the arm model were tested for with Student’s t-tests on the mean for each direction of reaching.

### Data Availability

The anonymised datasets generated during the current study are available from the corresponding author on reasonable request.

## Author contributions

AAF conceived the problem, SVT and AAF designed the experiments. SVT performed the experiments, SVT and AAF performed the modelling and analysis. SVT and AAF wrote the manuscript.

## Supplementary Information

**Figure S1.**
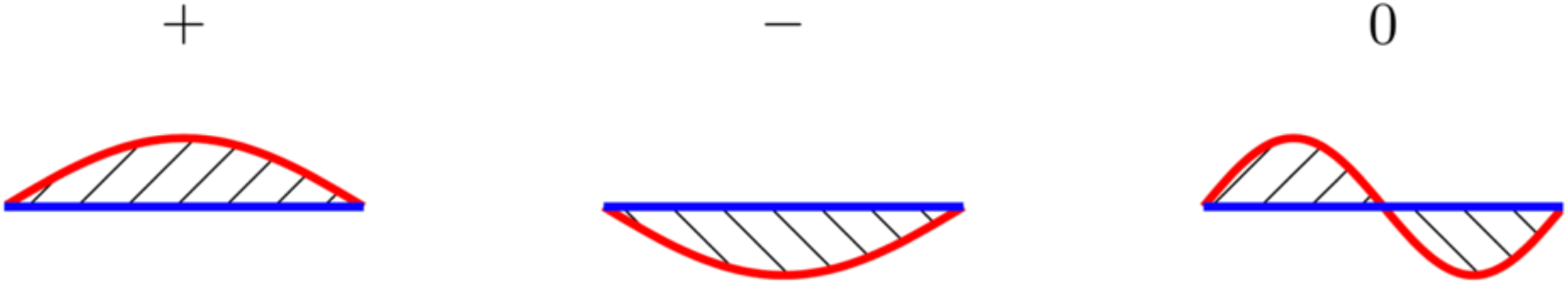
**Left**: Integrating the area between the red and blue trajectories produces a positive value. **Centre**: Here, integrating the area between the red and blue trajectories produces a negative value. **Right**: Finally, integrating the area between the red and blue trajectories produces 0 as the first and second halves of the traces cancel each other out.

**Figure S2.**
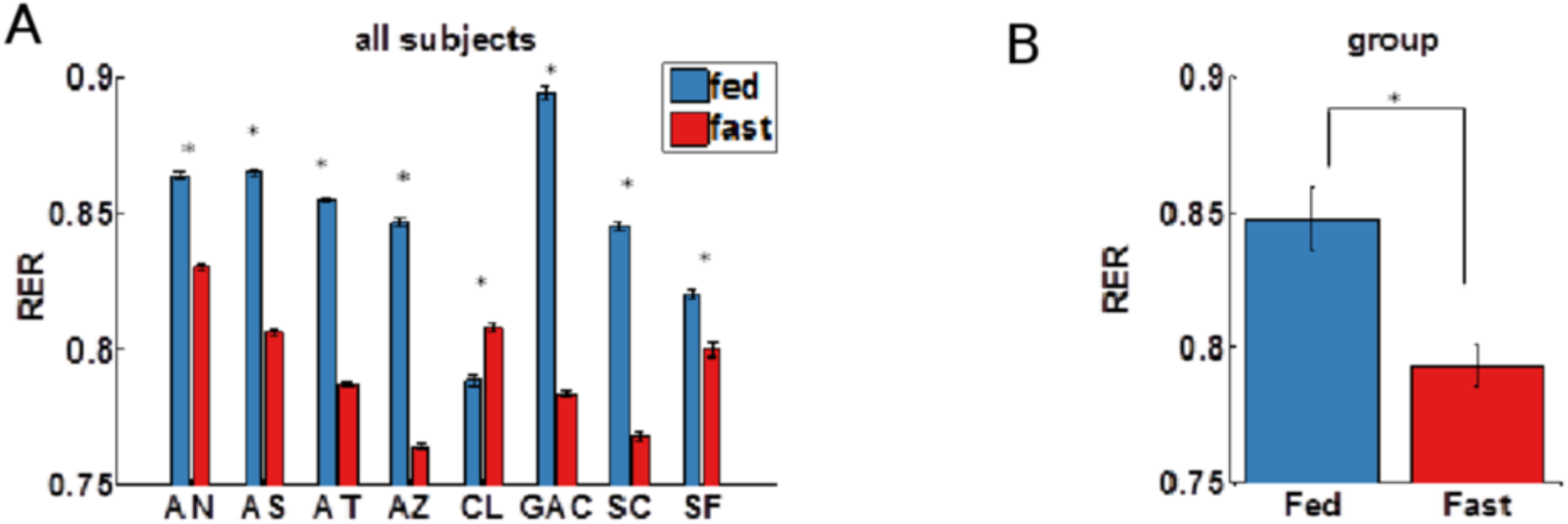
**(A)** Fed (blue) and fasted (red) respiratory exchange ratio is shown for each subject (labelled on the x axis), with standard error indicated by error bars. Significant differences are indicated with a *. A decrease in RER was expected when fasted and we observed this decrease in 7 out of 8 experimental Subjects (Student’s t-test, p < 0.00005 for each of the 7 subjects). **(B)** Fed (blue) and fasted (red) respiratory exchange ratio is shown for the group, with standard error indicated by error bars. The change in RER was significant for the group (Student’s t-test, p=0.0075), denoted with an asterisk.

**Figure S3.**
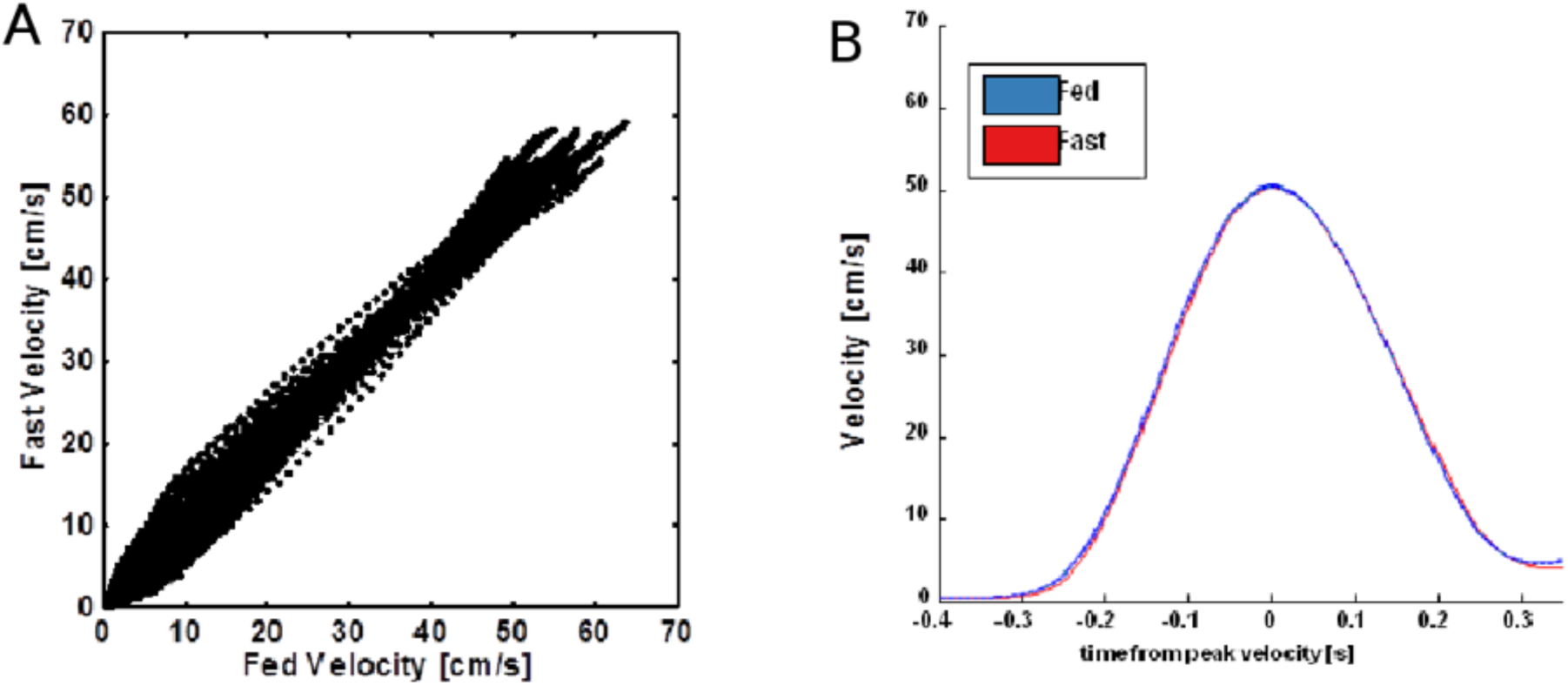
**(A)** Mean fasting velocities plotted versus mean fed velocities for all subject/direction combinations. Note the even distribution about the line of equality. **(B)** Mean velocity traces ± standard error plotted versus time from peak velocity for an exemplary direction. Fed (blue) and fasted (red) velocity traces are barely distinguishable and the standard error is too small to be easily visible on this plot.

